# Detection of *Plasmodium* infections in macaques from areas endemic for brugian filariasis in Belitung District, Indonesia

**DOI:** 10.64898/2026.08.10.743871

**Authors:** Irina Diekmann, Taniawati Supali, Elisa Iskandar, Matthew R. Kulpa, Noviani Sugianto, Rahmat Alfian, Yossi Destani, Abakar Gankpala, Kerstin Fischer, Balbir Singh, Paul C.S. Divis, Peter U. Fischer

## Abstract

**Background:** Malaria caused by *Plasmodium knowlesi* and lymphatic filariasis caused by *Brugia malayi* are mosquito-borne infections with non-human primates as reservoirs. *P. knowlesi* has emerged as a significant cause of human malaria in Southeast Asia over the past two decades. Belitung district, Indonesia, was until recently assumed to have eliminated *B. malayi* until infections were detected in humans and long-tailed macaques. To investigate whether the local reservoir of *B. malayi* is also a reservoir for malaria we screened macaques from 4 areas in Belitung for malaria parasites.

**Methods and findings:** Blood samples from 163 long-tailed macaques (*Macaca fascicularis*) that had been tested for *B. malayi* were examined by quantitative PCR assays specific for *Plasmodium* spp., *P. knowlesi*, *P. inui*, *P. coatneyi* and *P. cynomolgi*. A total of 130 macaques (79.8%) tested positive in the pan-*Plasmodium* qPCR assay. *Plasmodium inui* was most prevalent (41.7%), followed by *P. knowlesi* (38.7%), *P. coatneyi* (24.5%) and *P. cynomolgi* (13.5%). Multiple species infections, with 2-3 *Plasmodium* species were detected in 37% of macaques. Notably, 20 (91%) of 22 *B. malayi*-positive macaques were co-infected with at least one *Plasmodium* species. We sequenced the complete mitochondrion from 9 samples diagnosed by qPCR as mono-infections. Phylogenetic analysis confirmed 7 as *P*. *knowlesi*, and the other two as *P*. *inui* and *P*. *coatneyi*. Phylogenetic and pairwise analysis revealed that *P*. *knowlesi* isolates from Belitung were closely related to each other and to *P*. *knowlesi* from humans and monkeys from Thailand, Malaysia and Indonesia.

**Conclusions:** Molecular evidence shows high prevalence of zoonotic malaria parasites in macaques from Belitung, emphasizing the risk of human transmission. Increased surveillance, improved diagnostics, and targeted interventions are needed to prevent zoonotic spillover of *P. knowlesi* as it has been observed for *B. malayi* in Belitung.

**Author summary:** We examined macaques for *Plasmodium* parasites in Belitung Island, Indonesia, a region classified as free of locally transmitted human malaria. *Plasmodium knowlesi* is a monkey parasite that commonly infects humans and has emerged as a concern in Southwest Asia over the last 20 years. In Belitung, macaques are infected with *Brugia malayi,* a filarial nematode, that causes lymphatic filariasis in humans. Blood samples from 163 macaques were screened for *Plasmodium* DNA and 80% carried malaria parasites. *Plasmodium inui* was most common, followed by *P. knowlesi, P. coatneyi,* and *P. cynomolgi*. Many macaques were infected with multiple parasites simultaneously, but no host was infected with all four *Plasmodium* species. Ninety one percent of *B. malayi*-positive macaques were co-infected with malaria parasites, including *P*. *knowlesi*, indicating multi-parasite infections that could potentially spread to humans. Analysis of the mitochondrial genome of 7 *P. knowlesi* isolates from Belitung showed that they were most similar to each other and both human and monkey samples from Malaysia, Thailand and Indonesia. The study highlights the need for enhanced malaria monitoring and prevention, given the complex epidemiology of co-infecting parasites and risk to humans and animals.

## Introduction

*Plasmodium knowlesi* is a zoonotic parasite that has emerged as important cause of human malaria in Southwest Asia over the last twenty years [1]. The natural hosts of *P. knowlesi* are primarily long-tailed and pig-tailed macaques [2]. These macaques can host up to seven *Plasmodium* species: *P. knowlesi, P. cynomolgi, P. coatneyi, P. inui, P. inui-like, P. simiovale* and *P. fieldi* [3]. Although *P. knowlesi* is currently the most significant simian malaria parasite infecting humans, other zoonotic malaria cases have recently been reported in Southeast Asia. These cases were caused by *P. cynomolgi* and *P. inui* in Malaysia and Thailand, by *P. fieldi* in Thailand and by *P. coatneyi, P. inui-like and P. simiovale* in Malaysia [4–6]. The infections were detected using molecular methods, including nested and quantitative PCR assays, as well as DNA sequencing. These methods are more accurate and sensitive for the identification of malaria parasites compared with the examination of blood smears [7].

In Indonesia, 98% of human malaria cases are caused by *P. falciparum* and *P. vivax* that do not have an animal reservoir, but human cases due to *P. knowlesi* are increasing [8, 9]. Given the large size of Indonesia, there have been only very few studies that assessed non-human primates for zoonotic malaria. One study collected 108 long-tailed and pig-tailed macaques in Aceh, North Sumatra, West Sumatra, Central Java and Central Kalimantan and detected *Plasmodium* DNA in 51% of the individuals with 8% being *P. knowlesi* [10]. In another study that included 70 long-tailed macaques from Bintan and Sumatra islands one of them was infected with *P. knowlesi*, but 93% harboured *P. cynomolgi* or *P. inui* [11]. Taken together, these reports indicate on one hand a widespread distribution of macaques with zoonotic malaria parasites in Indonesia and a potential infection risk for humans, but also on the other hand a lack of comprehensive data because of small sample size and low geographic coverage within the large distribution area of macaques.

The Indonesian Ministry of Health has targeted malaria for elimination using a stepwise subnational approach. In this approach the Belitung and Belitung Timur districts in western Indonesia were classified to be free of human malaria [9, 12]. However, countries where transmission of “human malaria” has been interrupted but *P. knowlesi* cases persist in humans, cannot be officially certified by WHO as malaria-free [13]. In parallel Indonesia has joined the Global Program to Eliminate Lymphatic Filariasis (GPELF) of the WHO and in 2017 it was assumed that Belitung district had eliminated lymphatic filariasis. This neglected tropical disease is caused in eastern Indonesia mostly by *Brugia malayi.*. Post elimination surveillance of lymphatic filariasis showed that after 5 rounds of antifilarial mass drug administration active *B. malayi* infections in adults were still detected [14]. A subsequent survey examining the potential animal reservoir for *B. malayi* showed approximately 14% of 163 long-tailed macaques (*Macaca fascicularis*) were infected with *B. malayi* [15]. Whether the same macaques are also a reservoir for *P. knowlesi* and pose a risk not only for transmitting lymphatic filariasis but also malaria is not known. *Plasmodium knowlesi* is transmitted by *Anopheles* mosquitoes while nocturnally subperiodic *B. malayi* is usually transmitted by *Mansonia* species [16, 17]. However, laboratory studies have shown that some *Anopheles* species or selected strains do support the development of nocturnally periodic *B. malayi* [18]. Co-endemicity and potentially co-transmission by the same vectors or vector species with similar biting habits of zoonotic malaria and lymphatic filariasis would warrant integrated intervention for both diseases.

Genome sequencing and population genomic studies have revealed complex patterns of genetic diversity and host-association, along with unique demographics that distinguish *P. knowlesi* from other human malaria parasites [19]. For example, analysis of mitochondrial DNA (mtDNA) sequences revealed variations among *P. knowlesi* infections in different hosts with macaques typically showing more variable haplotypes per infection and higher overall diversity than those found in humans [20]. In addition, identical *P. knowlesi* mtDNA haplotypes were found in both humans and macaques, and no major type was associated exclusively with either host within the same geographical region, suggesting ongoing zoonotic transmission rather than recent host adaptation. Therefore, a further characterization of local *P. knowlesi* isolates by comprehensive mtDNA sequencing should be useful to build a data base.

Within the framework of an integrated assessment of macaques as animal reservoir for malaria and lymphatic filariasis the objective of the current study was to screen blood samples from macaques collected in 5 villages for *Plasmodium* DNA, *P. knowlesi* and 3 other zoonotic malaria parasites. We used a combination of established and novel qPCR assays for species identification and sequencing of the mitochondrial genome to better characterize the local *P. knowlesi* strains. We analysed co-infection of *Plasmodium* species and compared them with *B. malayi*. The results call for integrated one health research for malaria and lymphatic filariasis to expand our current understanding of potential public and wildlife health concerns.

## Materials and methods

### Ethics approval and consent

The trapping and blood collection of animals were approved by the Ministries of Health and the Environment and Forestry of Indonesia (protocol #22-040365). The study received ethical approval from the ethical committee of Universitas Indonesia (no 515/UN2.F1/ETIK/PPM.00.02/2022) and was performed by veterinarians or, under their supervision, by veterinary technicians.

### Study area and sample preparation

The study was conducted in Belitung Island, Indonesia, that is divided into two districts: Belitung and Belitung Timur. Blood samples were collected from wild macaques in five locations within the Belitung district: The neighbouring villages - Selat Nasik and Petaling on Mendanau Island, Kembiri, Lassar, and Kacang Butor on the main island of Belitung. The study area and sample collection have been described in detail in a previous publication. [15]. Briefly long-tailed macaques were trapped, and samples were collected either between 07:30-10:30 am or 3:00-7:30 pm when animals were most active. A total of 163 macaque blood samples were collected (Table S1) [15].

Giemsa stained three-line thick blood smears were examined for filarial parasite species in the field as described previously [15] but not for *Plasmodium* species. Aliquots of the blood samples were stored at -20°C and accurate morphological species identification of the intracellular *Plasmodium* parasites in the laboratory was not possible.

### DNA extraction and real-time PCR

DNA was extracted from 50 µL of each blood sample. Using the Qiagen Blood and Tissue Kit (Hilden, Germany), total DNA was obtained from 50 μL of animal blood (in EDTA) following the manufacturer’s instructions. The elution was performed with 200 μL DEPC-treated water. The extracted DNA was stored at -20°C until further use. Real-time PCR assays were performed to detect *Plasmodium spp*., *P. knowlesi, P. cynomolgi, P. coatneyi* and *P. inui.*

Real-time PCR was conducted using a QuantStudio 6 FLEX (Applied Biosystems - Thermo Fisher, Waltham, MA) Thermocycler with a total reaction volume of 10 µl. The reaction mixture contained 1 µl of DNA, 5 µl of TaqMan Fast Master Mix (Thermo Fisher), and 1 µl of primer/probe mix (Integrated DNA Technology, Coralville, IA). The cycling protocol involved an initial pre-read at 60 °C for 30 seconds, a hold at 95 °C for 20 seconds, followed by 40 cycles of 95 °C for 1 second and 60 °C for 20 seconds, and a final post-read at 60 °C for 30 seconds. Pan-*Plasmodium* primers used were published previously by Kamau and colleagues [21]. Positive samples were further investigated using *Plasmodium*-specific primer [22] in combination with species-specific probes published previously (Table S2) [23, 24]. Positive controls for the PCR were received for all four species from Malaysia, or for *P. knowlesi* and *P. cynomolgi*, DNA was provided by NIH/NIAID BEI Resource Center (www.beiresources.org). The samples were processed in duplicates. For samples with a high CT value and for those further processed for mtDNA amplification, the real-time PCR was repeated for verification. The presence of filarial nematode DNA, including *B. malayi*, was assessed by qPCR as described in a previous study [15].

### Mitochondrial genome assembly and phylogenetic analyses

The mtDNA was amplified using conserved primers described by Jongwutiwes et al. [25]. Primers were first evaluated using *Plasmodium* positive control before field samples were tested. DNA The standard PCR protocol involves a 50 μl reaction mixture containing 0.2 μM of each primer, and 1X of PrimerStar Taq™ DNA polymerase (Takara, Seta, Japan). The thermocycling protocol involved 35 cycles with an initial denaturation at 94°C for 60 seconds, denaturation step at 96°C for 20 seconds followed by annealing at 52°C for 15 seconds Extension step was at 72°C for 10 minutes. Sequences were manually assembled and aligned to *P. knowlesi* mtDNA and other relevant *Plasmodium* sequences using MEGA11 (https://www.megasoftware.net/). Annotations were performed via GeSeq [26] and all nine mitochondrial sequences were submitted to NCBI GenBank (Accession numbers: PZ347423- PZ347431; Table S3).

In order to perform phylogenetic analyses, mitochondrial sequences (n=43) were first aligned in Geneious Prime v2026.0.2. This included the nine unknown samples that were identified through BLAST analysis. Seven of which were identified as *P*. *knowlesi* and the other two were identified as *P*. *coatneyi* and *P*. *inui*. The two coinfections of *P. knowlesi* and *P. coatneyi* or *P.inui* were later confirmed by qPCR. Three sequences from *Leucocytozoon* species, blood-inhabiting protozoan of the same order, functioned as the phylogenetic outgroup. The alignment was then exported to raxml 2.0.16 [27] to perform a maximum likelihood analysis (ML) with 1000 bootstrap replications. Phylogenetic tree topology and formatting was assembled in FigTree v1.4.4 software (https://tree.bio.ed.ac.uk/software/figtree/). In addition, pairwise comparisons were done by calculating the genetic diversity within/among *P*. *knowlesi* sequences from this study and from GenBank using a maximum composite likelihood (Table S4).

## Results

### Detection of *Plasmodium* DNA by real-time PCR

Before testing macaque blood samples, plasmids comprising DNA from *P*. *knowlesi*, *P*. *inui*, and *P*. *cynomolgi* were used to assess pan-*Plasmodium* assay. The limit of detection for *Plasmodium* plasmid DNA for *P*. *knowlesi* and *P*. *inui* was approximately 1,000 ag/µL and 10 ag/µL for *P*. *cynomolgi.* Linear regression analyses of the cycle threshold (Ct) showed high dependence on DNA concentration for all three species (*P*. *knowlesi:* R² = 0.9656, *P*. *inui*: R² = 0.9609, and *P*. *cynomolgi*: R² = 0.9942) (Fig S1). When using the pan-*Plasmodium* assay, the average Ct value was 30.40 (range 23.74 – 38.35) for all positive samples. Average Ct values were relatively consistent across *Plasmodium* species with 30.14 (23.38 – 39.60) for *P*. *inui*, 29.82 (22.44 – 37.81) for *P*. *knowlesi*, 30.37 (24.89 – 36.54) for *P*. *coatneyi* and 31.67 (24.27 – 39.44) for *P*. *cynomolgi*. Among the 397 reactions with positive Ct values below 40, 39 samples (9.8%) had values between 33 and 35 and 46 samples (11.6%) had values >35. These results show that the qPCR assays used are highly sensitive and when applied to field samples, the majority of samples produced relatively low Ct values that allowed a clear differentiation of positive and negative samples.

Out of 163 macaques, 130 (79.8 %) tested positive for *Plasmodium* DNA using the pan-*Plasmodium* qPCR assay and each of these were positive for at least one *Plasmodium* species as detected by the species-specific assays (Table 1). The *Plasmodium* species that was most commonly detected was *P. inui* which was found in 41.7% of samples tested. The second most frequent infection detected was *P. knowlesi* (24.5%), followed by *P. coatneyi* (24.5 %), and *P. cynomolgi* (13.5%). Of the five villages that samples were collected from, the highest percentage of *B*. *malayi, P*. *knowlesi*, and *P*. *cynomolgi* infections occurred in macaques from Petaling village. Moreover, the highest percentage of *P*. *inui*, and *P*. *coatneyi* were found in macaques from the Lassar and Selat Nasik villages, respectively.

**Table 1.**
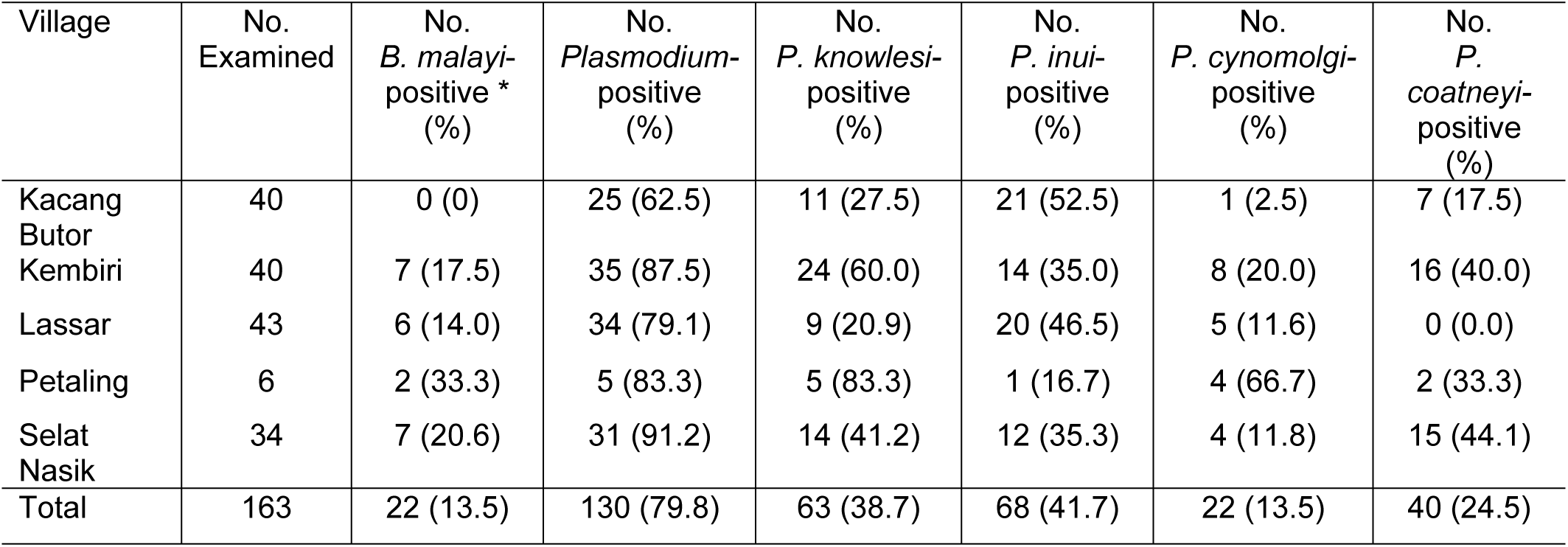
Detection of the blood parasites *B. malayi* and *Plasmodiun* spp. in macaques collected in various villages in Belitung District, Indonesia. * Data from [15]

### Analysis of co-infections

Co-infection with more than one species of *Plasmodium* occurred in 54 (33%) of the macaques; however, none of the samples confirmed infections with all four *Plasmodium* species (Table 2). Notably, among the 22 macaques that tested positive for *B. malayi* [15], 20 (91%) were co-infected with one or more *Plasmodium* species, with the highest co-infection observed with *P*. *knowlesi* followed by *P*. *inui*, *P*. *coatneyi*, and *P*. *cynomolgi* (Table 2). Overall, 79.8% of the macaques were *Plasmodium* positive and 91% of *B. malayi* positive macaques were also *Plasmodium* positive, indicating that macaques infected with *B. malayi* are more likely to be also infected with *Plasmodium*.

**Table 2.**
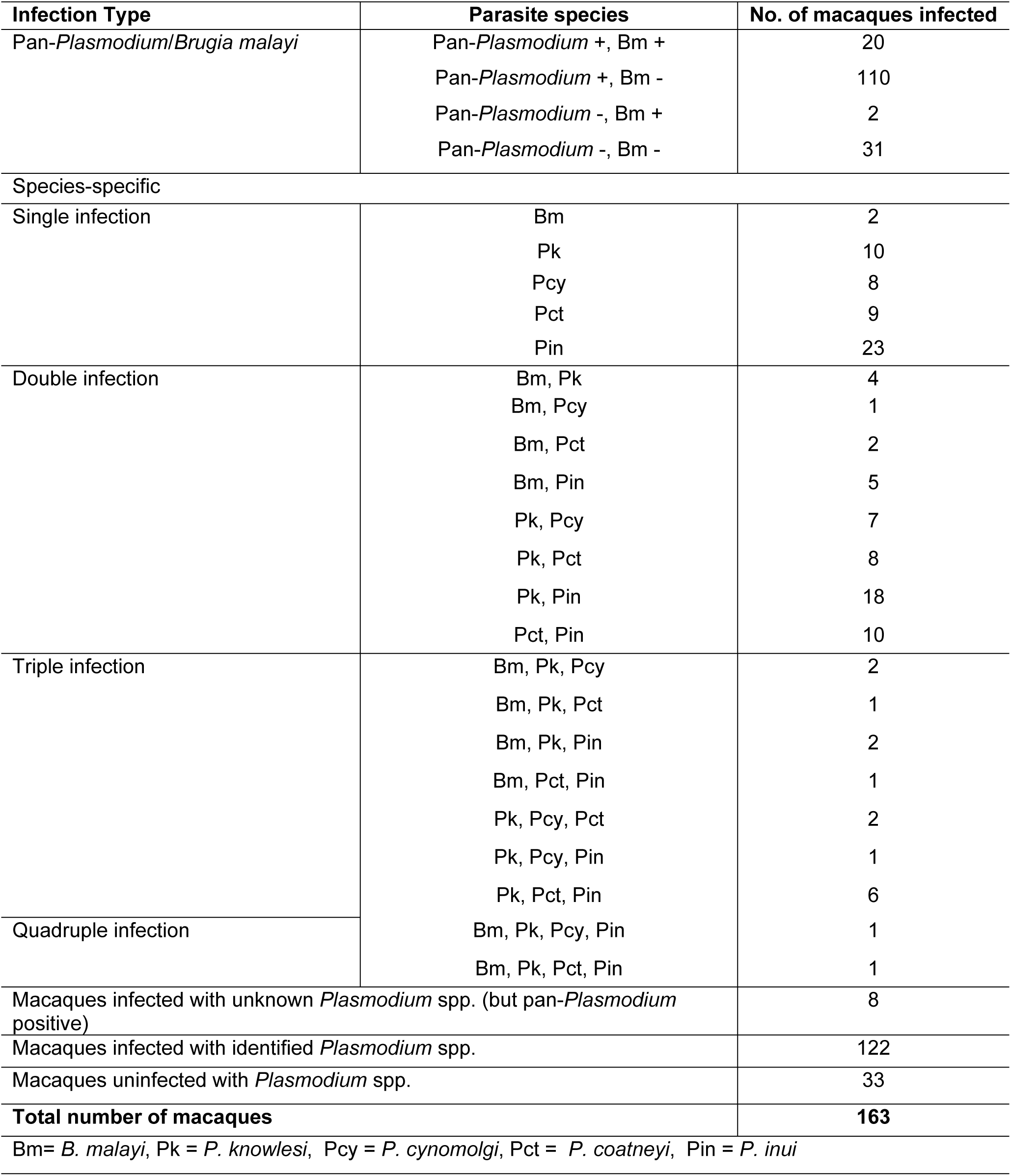
Detection of co-infecting parasites, *B. malayi* and *Plasmodium* spp., in macaques collected in Belitung District, Indonesia. Multiple infections that were not detected in the 163 tested macaques are not shown.

### Mitochondrial genome analysis

Phylogenetic analysis of mitochondrial sequences from *Plasmodium* species in macaque blood (n=9) and NCBI GenBank (n=31) formed three distinct clades with 100% bootstrap support. Two of these clades comprised only a single *Plasmodium* species (i.e., *P*. *falciparum* and *P*. *gallinaceum*). The third clade comprised all other remaining species that were used in analysis including *P*. *coatneyi*, *P*. *cynomolgi*, *P*. *fieldi*, *P*. *inui*, *P*. *knowlesi*, *P*. *simiovale*, and *P. vivax*. All *P. knowlesi* from macaque blood (n=7) and NCBI GenBank (n=14) formed a single monophyletic clade with 100% bootstrap support. This topology was found similarly in other *Plasmodium* species sequences that were analyzed as well (Fig 1).

**Fig 1.**
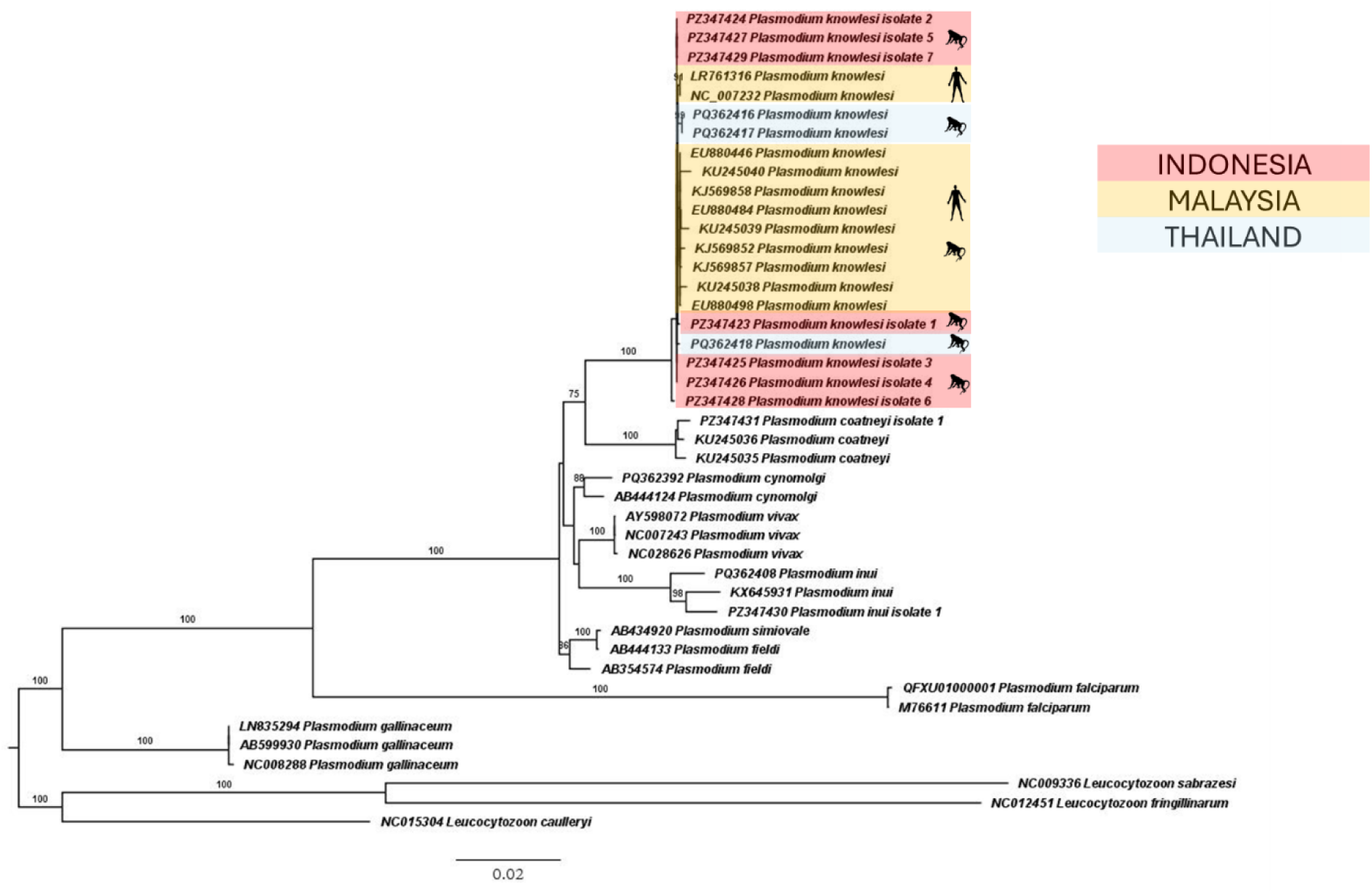
Phylogenetic tree based on mtDNA of *Plasmodium* and *Leucocytozoon* spp using a maximum likelihood approach. Figures on branches are bootstrap percentages based on 1000 replicates and only those above 74 are shown. Genbank Accession numbers are shown for each isolate.

In addition, 48 *P*. *knowlesi* mitochondrial sequences from NCBI GenBank were used to make pairwise comparisons with our recently uncovered *P*. *knowlesi* sequences (n=7). Overall, *P*. *knowlesi* sequences from Belitung had on average more similarity to *P*. *knowlesi* sequences originating from macaques in southern Thailand (n=3; 0.11% (0.06-0.19%)) and least similarity to *P*. *knowlesi* sequences originating from macaques in Sarawak, Malaysia (n=4; 0.28% (0.17-0.49%)). There was little pairwise difference when comparing our *P*. *knowlesi* sequences between others infecting macaque (n=30; 0.15% (0.06-0.49%)) versus human hosts (n=18; 0.14% (0.06-0.27%)). Intra-isolate comparisons were also analyzed and revealed an average of 0.05% (0.00-0.14%) genetic distance between sequences (Table S4).

## Discussion

In this study, blood samples from 163 long-tailed macaques were examined for *Plasmodium* spp. These samples were collected as part of a broader investigation into animal reservoirs of *B. malayi* in Belitung Island, Indonesia[15]. While in recent years non-human primates from many regions in southeast Asia have been screened for *Plasmodium* [28–31], no studies have simultaneously tested macaques for both parasites that can cause malaria and lymphatic filariasis in humans. Nearly 80% of the samples collected from macaques in Belitung were infected with *Plasmodium* and almost half of these infections were positive for *P. knowlesi*. Prior to this study, Belitung Island was considered a non-endemic area for human malaria infections [9, 12]. Only sporadically human malaria cases were reported by district hospitals or primary health care centres (Puskesmas) to the health authorities of Belitung and Belitung Timur districts (TS personal communication). These malaria cases were not further specified and assumed to be imported from close by Bangka Island or other endemic areas based on travel history. The present study demonstrates that in Belitung, macaques are a natural reservoir for two major parasitic diseases targeted for elimination in Indonesia, malaria and lymphatic filariasis.

The real-time qPCR assays used in this study were designed to detect four of the five *Plasmodium* species commonly detected in macaques, namely *P. knowlesi, P*. *coatneyi*, *P*. *cynomolgi*, and *P*. *inui*. A specific probe for the fifth species, *P. fieldi,* and for the more recently described *P. simiovale* and *P. inui-like* parasites were not available at the time of analysis. Therefore, potential cross-reactivity with closely related species or the presence of undetectable co-infections was not tested. Samples comprising only *P. knowlesi* infection were identified in seven samples and confirmed by sequencing of the complete mtDNA. Although the qPCR was run in repeat, two samples that were assumed to be *P. knowlesi*-only infections were either cross-reactive or, more likely, had co-infections with another *Plasmodium* species. Sequencing of the mtDNA revealed *P*. *coatneyi* and *P*. *inui*. Because only a single mitochondrial genome was assembled per sample and primer walking was sequenced based and not species based, it is possible that the conserved PCR products were not from *P. knowlesi.* The probes we used were originally designed based on Malaysian reference sequences, and genetic differences in Indonesian strains may reduce probe binding efficiency. Such sequence mismatches could potentially cause decreased sensitivity or false-negative results. Also, about 5% of macaque samples were pan-*Plasmodium* qPCR positive but could not be further specified. Since probes for *P. fieldi, P. simiovale* and *P. inui*-like species were not available during the current study, it is likely that a larger variety of *Plasmodium* infections in macaques occurs

Previous molecular diagnostic studies to detect zoonotic malaria usually used reverse-transcriptase qPCR, nested or hemi-nested PCR assays [32–35]. Our results show that a single step qPCR approach is a convenient screening tool to detect *Plasmodium* parasites in non-human primates. Compared to nested and hemi-nested PCR, the qPCR assays are faster and less prone to contamination because no handling of amplified DNA is necessary. In areas where suspected human malaria cases are rarely reported, like in Belitung, diagnosis of malaria based on clinical features is difficult because of non-specific and overlapping clinical signs with many other infectious diseases. Even if blood smears are examined by microscopy, accurate morphological diagnosis of zoonotic malaria by light microscopy is challenging since simian malaria parasites share similar morphological characteristics with human malarias: *P. knowlesi* and *P. inui* are both similar to *P. malariae, P. cynomolgi* is identical to *P. vivax* and *P. coatneyi* resembles *P. falciparum* [36], [37]. Therefore, improved molecular diagnosis for *P. knowlesi* and other zoonotic malaria parasites is essential to determine whether humans in Belitung are infected.

Phylogenetic analysis of the complete mtDNA of *P. knowlesi*, including reference sequences compiled from NCBI GenBank, revealed that they were closely related to each other and formed a single monophyletic clade (Fig. 1). The close phylogenetic relationship to the Malaysian samples infecting humans suggests a potential zoonotic origin of the Belitung *P. knowlesi* strain. In addition, *P*. *coatneyi*, a *Plasmodium* species of simian primates only recently linked by molecular studies to a few human infections [6], is most closely related to *P*. *knowlesi*. Similar studies from southern Thailand also show a close relationship between *P. knowlesi* and *P. coatneyi* [29]. Given this close relationship and the reported human cases *P*. *coatneyi* should be monitored more closely. Pairwise analysis of mtDNA revealed *P*. *knowlesi* isolates from Belitung and other locations were all closely related. However, in this study *P*. *knowlesi* from macaques were most genetically similar to other *P*. *knowlesi* infecting macaques from Thailand. This could imply that the subpopulation of *P*. *knowlesi* established on Belitung Island and the Sumatra region were translocated from regions of Thailand or mainland Malaysia (or vice versa) rather than peninsular Malaysia or other parts of Borneo. Previous population genomic studies identified distinct subpopulations of *P. knowlesi* in mainland and Borneo [38, 39]. However, more data is needed to support these assertions. The analysis of the mitochondrial genome showed little difference between the isolates recovered from Belitung compared to *P*. *knowlesi* strains infecting either humans or macaques, independent of geographic region. This suggests that there is only a single strain of *P*. *knowlesi* in Belitung that, while it is found in macaques, it is equally capable of infecting human hosts (Table S1). More surveillance testing of humans and macaques is required to understand *P*. *knowlesi* strain host associations of Belitung Island.

Macaques are the natural reservoir for *P*. *knowlesi* and are often simultaneously infected with several other simian malaria parasites [29, 31, 35, 40]. In our study, mixed-species *Plasmodium* infections in macaques were more common than single-species infections that varied between 10% for *P. coatneyi* and 41% for *P. cynomolgi*. More than 90% of macaques that were infected with *B. malayi* were co-infected with at least one *Plasmodium* species and 55% of them were co-infected with *P. knowlesi*. Potential co-infections of humans with the related filarial parasite *Wuchereria bancrofti* and *P. knowlesi* have been discussed previously with regard to integrated control and surveillance [16]. However, our study reports the first co-infections of *B. malayi* and *P. knowlesi* in a non-human primate reservoir.

Molecular xenomonitoring is recommended for surveillance of lymphatic filariasis, including *B. malayi* infection [41]. Filarial DNA can be detected in both vector and non-vector mosquitoes after a blood meal on infected hosts, but prevalence and amount of detected *B. malayi* DNA is higher in vectors [42]. Molecular xenomonitoring of *Plasmodium* infection and potential drug resistance is an important part of malaria surveillance [43, 44]. Although *P. knowlesi* and *B. malayi* may not share the same vector species, the high prevalence of co-infection in macaques indicate similar host preference of vector mosquitoes. Combining xenomonitoring efforts to include multiple pathogen agents, rather than focusing solely on a single pathogen, can reduce sampling effort and provide broader insights into the biology and epidemiology of future parasites of public health concern.

Only a limited number of studies assessed the prevalence of malaria parasites in wild and semi-captive macaques in Indonesia. These studies mainly focused on areas in parts of Sumatra, Kalimantan and Java. Overall, *Plasmodium* infection in macaques was common, with *P. inui* usually being the most prevalent species [10], [11] [45]. Even in a breeding facility in Bogor, Western Java, 14% of macaques tested positive for *P. inui* [46]. Because of the paucity of data from Indonesia a computer modelling study predicted a close to zero transmission suitability for *P. knowlesi* in Belitung in 2015 [47]. A revised classification by the same group in 2020 predicted an average transmission suitability of about 70% [48]. Our study provides clear evidence that Belitung is suitable for transmission of *P. knowlesi* with infection rates of macaques ranging between 21 and 60%.

In conclusion this study detected *P. knowlesi* and other *Plasmodium* infections in Indonesian macaques in Belitung, a non-endemic area for human malaria. It underscores the widespread distribution of zoonotic malaria parasites in non-human primates in Indonesia. Co-infections with *B. malayi* were common and may offer the opportunity of integrated surveillance. Further research is needed to investigate vector competence, interactions with other parasite species, and host adaptation in non-human primates and humans. Enhanced human surveillance and targeted public health interventions, including utilization of molecular detection methods, are essential to prevent misdiagnoses and avoid a potential increase in zoonotic transmission through targeted prevention measures.

## Acknowledgements

We thank Mr. Sudirman and the rest of the field team of the Universitas Indonesia for their expert technical support. Positive control DNA samples for *P. knowlesi* (plasmid, genomic DNA), *P. inui* (plasmid) and *P. cynomolgi* (plasmid) were provided by BEI Resources, NIAID, NIH.

## Ethics declarations

Trapping and blood collection of macaques were approved by the Ministries of Health and the Environment and Forestry of Indonesia (protocol #22-040365). The project received ethical approval from the ethical committee of Universitas Indonesia (no 515/UN2.F1/ETIK/PPM.00.02/2022).

## Consent for publication

Not applicable.

## Supporting Information

Figure S1 Dilution series and regression analysis for real-time qPCR assays detecting plasmid DNA with *P. knowlesi*, *P. inui* and *P. cynomolgi*.

Table S1 Individual collection and co-infection data of the examined macaques

Table S2 Sequences of primer and probes used in the present study

Table S3 GenBank accession numbers of DNA sequences used to generate Figure 2.

Table S4 Pairwise analysis of *P. knowlesi* sequences

